# Bacterial sensor evolved by decreasing complexity

**DOI:** 10.1101/2024.05.17.594639

**Authors:** Elizabet Monteagudo-Cascales, José A. Gavira, Jiawei Xing, Félix Velando, Miguel A. Matilla, Igor B. Zhulin, Tino Krell

## Abstract

Bacterial receptors feed into multiple signal transduction pathways that regulate a variety of cellular processes including gene expression, second messenger levels and motility. Receptors are typically activated by signal binding to ligand binding domains (LBD). Cache domains are omnipresent LBDs found in bacteria, archaea, and eukaryotes, including humans. They form the predominant family of extracytosolic bacterial LBDs and were identified in all major receptor types. Cache domains are composed of either a single (sCache) or a double (dCache) structural module. The functional relevance of bimodular LBDs remains poorly understood. Here, we identify the PacF chemoreceptor in the phytopathogen *Pectobacterium atrosepticum* that recognizes formate at the membrane distal module of its dCache domain, triggering chemoattraction. We further demonstrate that a family of formate-specific sCache domains has evolved from a dCache domain, exemplified by PacF, by losing the membrane proximal module. By solving high-resolution structures of two family members in complex with formate, we show that the molecular basis for formate binding at sCache and dCache domains is highly similar, despite their low sequence identity. The apparent loss of the membrane proximal module may be related to the observation that dCache domains bind ligands typically at the membrane distal module, whereas the membrane proximal module is not involved in signal sensing. This work advances our understanding of signal sensing in bacterial receptors and suggests that evolution by reducing complexity may be a common trend shaping their diversity.

**Significance:** Many bacterial receptors contain multi-modular sensing domains indicative of complex sensory processes. The presence of more than one sensing module likely permits the integration of multiple signals, although, the molecular detail and functional relevance for these complex sensors remain poorly understood. Bimodular sensory domains are likely to have arisen from the fusion or duplication of monomodular domains. Evolution by increasing complexity is generally believed to be a dominant force. Here we reveal the opposite – how a monomodular sensing domain has evolved from a bimodular one. Our findings will thus motivate research to establish whether evolution by decreasing complexity is typical of other sensory domains.

## Introduction

The ability of bacteria to sense and adapt to environmental changes is crucial for their survival. Bacteria have evolved an array of different signal transduction systems. Although different in architecture and mechanism, all signal transduction systems contain input and output modules (1). The canonical mechanism of action of signal transduction systems involves signal recognition at the input module, which is usually represented by a ligand binding domain (LBD), and signal transduction to modulate the output module. Hundreds of different LBDs have been described in bacterial receptors (2, 3) and new domains are discovered regularly (4, 5).

LBDs that detect signals in the extracytoplasmic space are typically flanked by transmembrane regions and are found in all major receptor types such as chemoreceptors, sensor histidine kinases, adenylate, diadenylate, and diguanylate cyclases; cAMP, -di-AMP, and c-di-GMP phosphodiesterases; and serine/threonine protein kinases and phosphatases (6, 7). Members of the same LBD family are frequently found in many different receptor types, indicating that LBDs recombine with different signaling domains to evolve proteins with different sensor functionalities (8–10). Although sequence-based classification of LBDs, such as that in the Pfam database (11), delineates many LBD types (2, 3), most of them belong to two structural folds, namely the α/β PAS/Cache fold and the four-helix bundle fold (3), each containing mono- and bimodular members (Fig. 1).

**Fig. 1.**
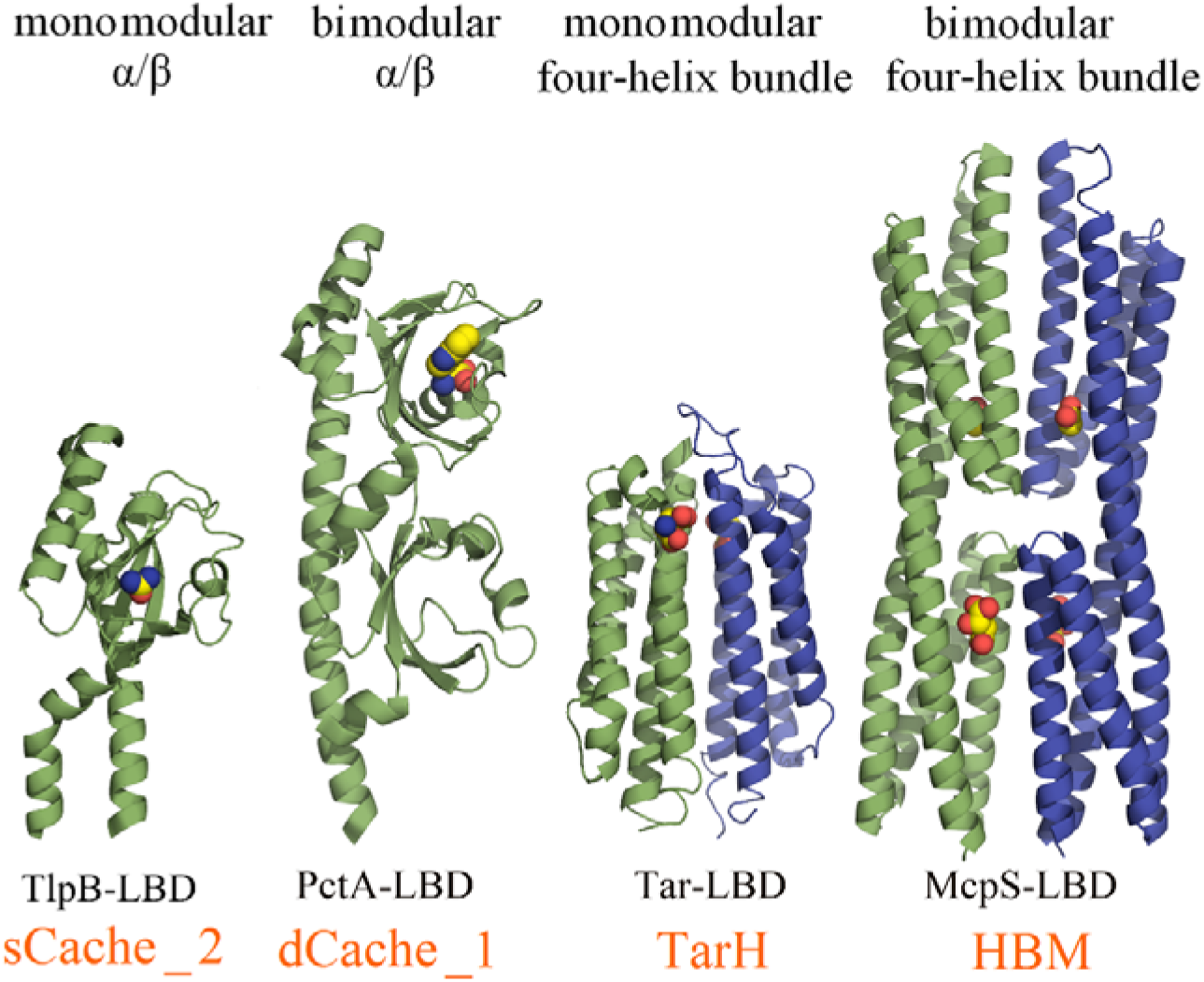
Mono- and bimodular ligand binding domains. Representative examples of the monomodular and bimodular bacterial ligand-binding domains. TlpB-LBD in complex with urea (sCache_2)(71), PctA-LBD in complex with L-Trp (dCache_1) (44), Tar-LBD in complex with L-Asp (TarH) (72), McpS-LBD in complex with malate and acetate (HBM) (14). Bound signals are shown in space-filling mode. Monomers of dimeric LBDs are shown in different colors.

Cache domains comprise the predominant superfamily of bacterial extracytosolic sensor domains and are found in all types of bacterial receptors (12). Cache domain-containing receptors are omnipresent in bacteria and have also been identified in archaea and many eukaryotes including humans (8). The bimodular dCache domains are likely to have arisen by duplication or fusion of monomodular sCache domains (12). The four-helix bundle type LBD is the most abundant LBD in chemoreceptors (13) and it is widespread in bacteria and archaea. Similarly to Cache domains, a bimodular version comprising two stacked four-helix bundles has been identified, termed the HBM domain (14) (Fig. 1).

The functional relevance of bimodular LBDs is poorly understood. One possible explanation is that different ligands bind to the individual modules resulting in an expansion of the sensing repertoire. This notion is supported by the demonstration that both modules of the bimodular HBM domain of the McpS chemoreceptor bind ligands (Fig. 1) and that both binding events trigger chemoattraction in an additive manner (14). On the other hand, the very large majority of bimodular dCache domains bind their ligands at membrane-distal module (15–22), whereas the membrane-proximal module remains unoccupied. Only few studies report signal recognition at both modules of dCache domains (23, 24).

dCache domains are classified in six families (12). The dCache_1 family (Pfam02743) is the most abundant and best characterized family. In contrast, no information is available for the Cache_3-Cache_2 domain (Pfam17201). This domain is likely to have arisen as result of a fusion of two mono-modular LBDs, namely sCache_3 and sCache_2 (12). Modelling indicates that this domain is composed of two α/β type modules linked by a long helix (Fig. S1); a structure similar to that of dCache_1 domains (25). Cache_3-Cache_2 domains were identified in chemoreceptors, histidine sensor kinases, diguanylate cyclases and phosphodiesterases (12). The initial objective of this study was to reveal the sensory capabilities of this domain and the functional role of Cache_3-Cache_2 domain-containing receptors.

Plant pathogens possess on average 27 chemoreceptors, which is nearly twice as many as the bacterial average of 14 (13), indicating that chemotaxis to diverse signals may be particularly important for bacteria that infect plants. This notion is supported by many studies showing that the deletion of chemoreceptors or chemosensory signaling genes reduces bacterial virulence (26–29). The interference with chemotactic signaling represents an alternative strategy to fight phytopathogens (30). However, there is only scarce information available on the signals that are recognized by phytopathogen chemoreceptors.

We are using *P. atrosepticum* SCRI1043 as a model strain to study chemoreceptor function. *P. atrosepticum* is among the 10 most relevant plant pathogens (31) and the causative agent of soft rot diseases (32). It has a single chemosensory pathway that contains 36 chemoreceptors. Only four of them have been characterized so far, responding to quaternary amines, amino acids and nitrate (15, 33, 34). SCRI1043 has one chemoreceptor, ECA_RS17860, with the Cache_3-Cache_2 domain (25).

We report here that it is a formate-specific chemoreceptor. We show that its bimodular Cache_3-Cache_2 represents an ancestral form from which monomodular sCache domains have arisen that preserved the capacity and molecular basis to bind formate. Bimodular Cache domains appear to have originated from monomodular domains by increasing structural complexity. This is the first report on a bacterial sensor that evolved by decreasing complexity. Further research will reveal whether similar events have also occurred in other families of bimodular sensory domains.

## Results

### The Cache_3-Cache_2 domain of ECA_RS17860 binds a single ligand, formate

To identify the function of chemoreceptor ECA_RS17860, we have generated its LBD as a purified individual domain and have conducted thermal shift-based ligand screening. This assay monitors ligand-induced increases in the midpoint of the protein unfolding transition (Tm) and Tm increase of 3 °C are considered a stringent threshold for significant binding. The screening of Biolog compound arrays PM1, PM2A, PM3B, PM4A, PM5, PM6, PM7 and PM8, each comprising 95 compounds, resulted in the identification of formate as a single ligand that significantly shifted the Tm (Fig. 2A). ECA_RS17860-LBD was then submitted to microcalorimetric titrations with Na formate (Fig. 2B), resulting in a dissociation constant of 18 ± 1 µM (Table 1), which is within the range affinities that is typically observed for ligand-LBD interactions (36). Chemoreceptor ECA_RS17860 was renamed PacF (*<u>P</u>ectobacterium <u>a</u>trosepticum* <u>c</u>hemoreceptor for <u>f</u>ormate).

**Fig. 2.**
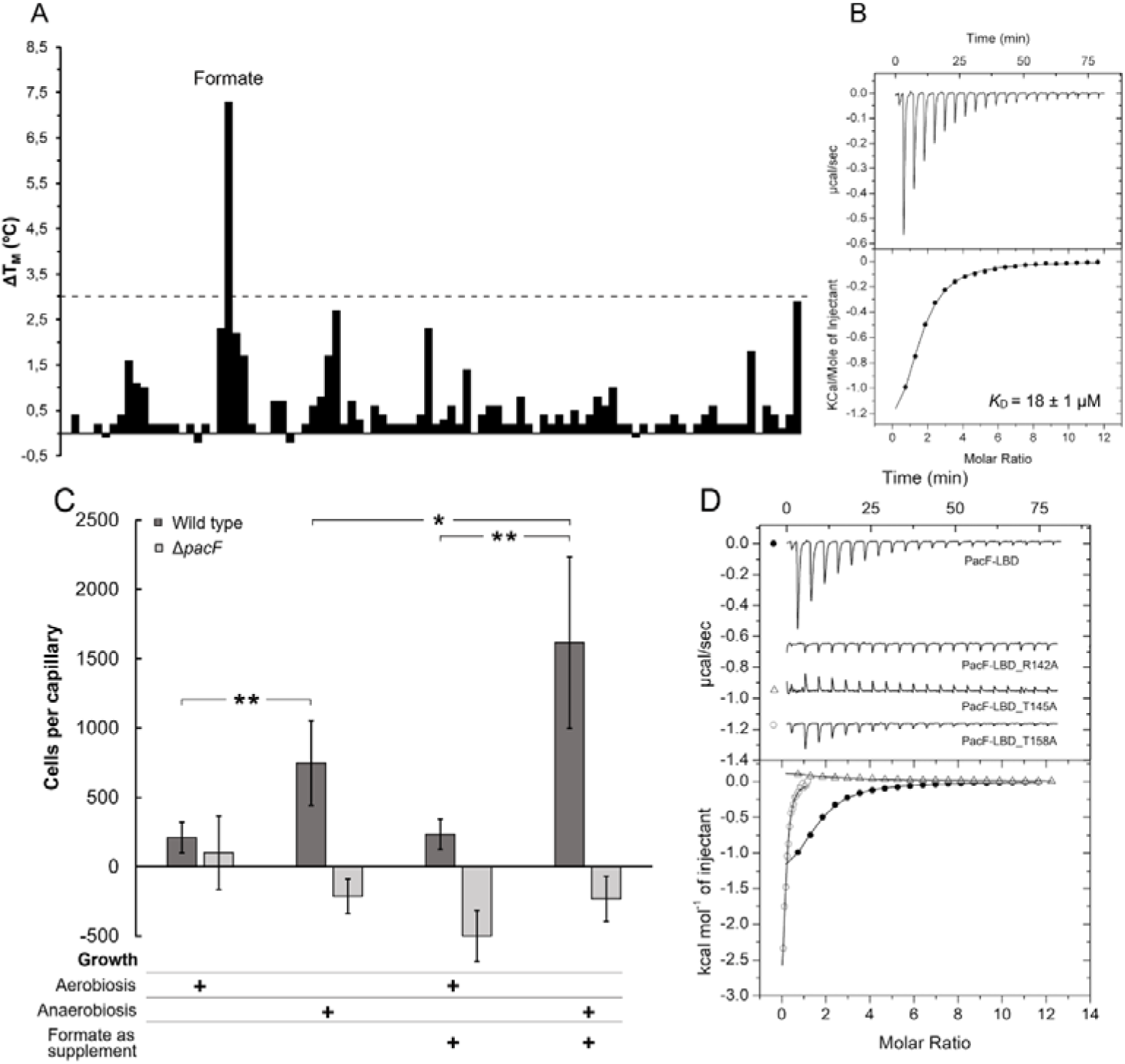
ECA_RS17860 (PacF) is a formate specific chemoreceptor. **A, B)** Formate binding to PacF-LBD. **A)** Thermal shift assays. Changes in the midpoint of the protein unfolding (Tm) caused by compounds of the PM1 array with respect to the ligand-free protein. The dashed red line indicates the threshold of 3 °C for significant hits. **B)** Microcalorimetric titration of 30 μM PacF-LBD with 5 mM Na formate. **C)** Quantitative capillary chemotaxis assays of *P. atrosepticum* and a mutant deficient in *pacF* to 0.1 mM Na formate. Cells were grown under aerobic or anaerobic conditions in the presence or absence of 0.5 mM Na formate. Data have been corrected with the number of bacteria that swam into buffer containing capillaries. *p < 0.05 in Student’s T- test; **p < 0.01 in Student’s T-test. D) Microcalorimetric titrations of 30 μM PacF-LBD and site-directed mutants with 5 mM (wt, R142A, T145A) or 500 μM (T158A) Na formate. **B, D**) Upper panels: Titration raw data. Lower panels: Integrated, dilution heat-corrected, and concentration-normalized peak areas fitted with the one-binding-site model of ORIGIN.

**Table 1.**
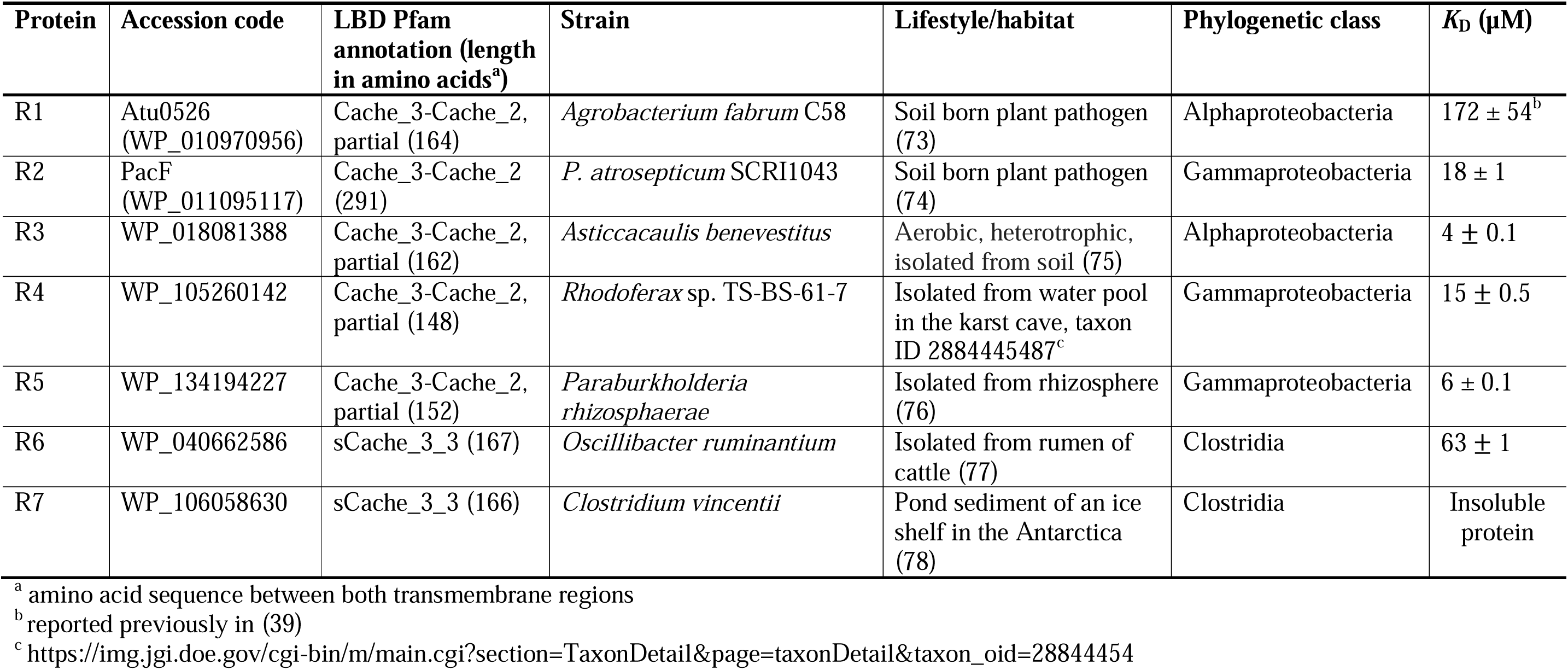
Microcalorimetric binding studies of Na formate to LBDs of different chemoreceptors. Indicated are the derived dissociation constants.

### PacF mediates chemoattraction to formate

To determine whether this chemoreceptor mediates chemotaxis in response to formate, we have constructed a deletion mutant that was together with the wild type strain subjected to quantitative capillary chemotaxis assays. Because *ECA_RS17860* transcript levels are significantly increased under anaerobic conditions (35), we have conducted chemotaxis assays with cultures grown in both aerobic and anaerobic conditions. Chemoreceptor transcript levels and the strength of chemotactic responses are frequently increased by the cognate chemoeffector(s) (36, 37). We have therefore assessed the effect of Na formate in the culture medium on the chemotactic responses. Under aerobic growth conditions weak responses to formate were observed; however, significantly stronger responses were seen when cells were cultured anaerobically (Fig. 2C, note: although cells were grown under anaerobic conditions, the chemotaxis assays were conducted under aerobic conditions). No chemoattraction to formate was observed in the chemoreceptor mutant under any experimental condition, indicating that PacF is the receptor that mediates chemotaxis towards formate. *P. atrosepticum* SCRI1043 was shown to perform formate respiration under anaerobic conditions (38). We conducted growth experiments showing that this strain is unable to use formate as carbon source for aerobic growth (Fig. S2). Formate is not toxic to SCRI1043 cells as evidenced by a minimal inhibitory concentration of above 50 mM.

### The membrane distal module of PacF-LBD is homologous to the single-module LBD of a formate chemoreceptor from *Agrobacterium fabrum*

Chemoreceptor Atu0526 from another plant pathogen, *A. fabrum* C58, has been reported to bind formate (39). Similarly to PacF-LBD, Atu0526-LBD matches the PfamCache_3-Cache_2 domain profile hidden Markov model (HMM) (11). However, the match is only partial, and Atu0526-LBD is much smaller than PacF-LBD (168 and 291 amino acid residues, correspondingly). It was shown that Atu0526-LBD binds formate directly and that the deletion of the *atu0526* gene abolishes formate chemotaxis (39). Modeling using Alphafold (40) showed that PacF-LBD is a bimodular dCache domain, whereas Atu0526-LBD is a monomodular sCache domain (Fig. S1). The sequence of Atu0526-LBD aligned well with the membrane-distal module of PacF-LBD (Fig. S3, Table S1) and both matched the same part of the Cache_3-Cache_2 HMM, suggesting that Atu0526-LBD is related to the membrane-distal module of PacF-LBD.

To confirm the formate binding site at PacF, we have generated three site-directed alanine replacement mutants of PacF-LBD in amino acid residues located in the membrane distal binding pocket (Fig. S1). R115 of Atu0526 was reported to be essential for formate binding (39). This residue is conserved in PacF (Fig. S3, R142) and its replacement also abolished formate binding as evidenced by microcalorimetric titrations (Fig. 2D). The mutation of two other binding pocket residues, that are also conserved in Atu0526 (Fig. S3), altered formate binding parameters in a differential manner. While PacF-LBD_T145A bound formate with an about 5-fold lower affinity (*K*_D_= 97 ± 15 µM), mutant T158A recognized formate with a 6-fold increased affinity (*K*_D_= 3 ± 0.3 µM, Fig. 2D).

### PacF-LBD is an ancestral domain from which the monomodular formate binding domain has evolved

We collected and analyzed a non-redundant set of 744 PacF-LBD homologs from various bacterial phyla (See Materials and Methods; Data S1). Whereas 164 of these sequences contained a sCache domain similar to Atu0526 (with a partial match to Cache_3-Cache_2 or a full-length match to sCache_3_3 HMM), the remaining 580 sequences had dCache domains similar to PacF (with a full-length match to Cache_3-Cache_2 HMM; Fig. 3A, Data S1). Significantly, the putative formate-binding residues were highly conserved in the entire dataset: R142 at 99% identity, and T145 and T158 at 98% identity (PacF residue numbering), suggesting that all homologs are formate binding LBDs.

**Fig. 3.**
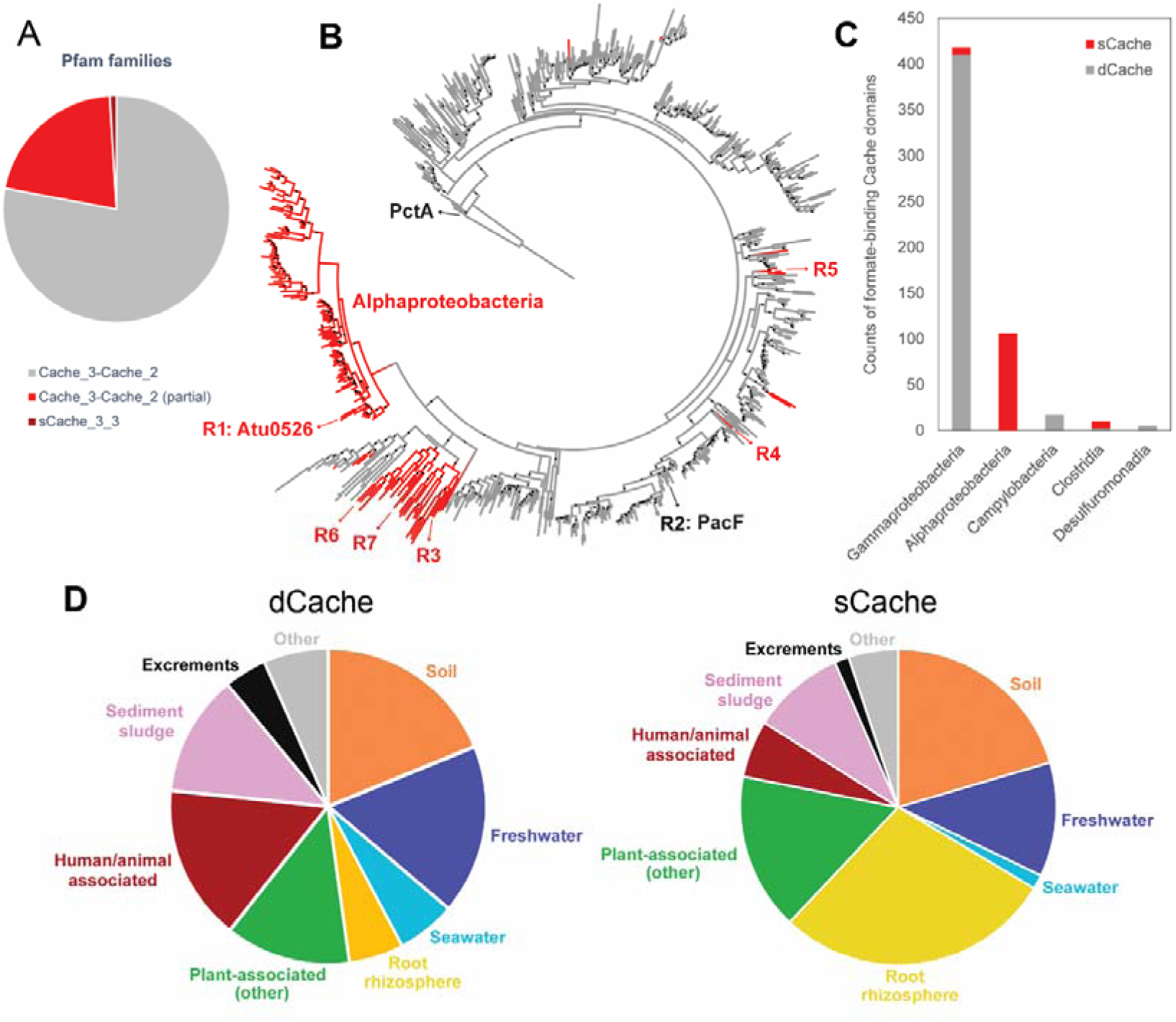
Formate-binding sCache and dCache containing LBDs. **A**) Pfam families of formate sensors. **B)** A maximum likelihood tree for formate-binding Cache domains (using membrane proximal modules of the dCache domain) built using MEGA with the JTT model and 100 bootstraps. Dots show branches with at least 70 bootstrap support. Red, sCache; grey, dCache distal module. The distal module of PctA dCache was used as the outgroup. Information on receptors R1 to R7 is provided in Table 1. The complete list of sequences is provided in Data S1. **C)** Distribution of formate sensors across bacterial classes based on the GTDB database. Classes with fewer than five proteins were not shown for simplicity. **D)** Isolation sources of strains containing dCache and sCache containing formate-responsive chemoreceptors.

We further built a maximum likelihood tree using the LBDs from these proteins (See Materials and Methods; Fig. 3B), which suggested that monomodular formate-binding sCache domains originated from the bimodular Cache_3-Cache_2 domain in several independent evolutionary events. This conclusion is further supported by the fact that the formate-binding sCache sequences are more similar to Cache_3-Cache_2 sequences than to other types of sCache domains. Protein sequences containing formate-binding Cache domains come from 571 genomes that are annotated in the GTDB database (41), and their distribution in bacterial classes is shown in Fig. 3C. Intriguingly, most formate sensors in gammaproteobacteria are dCache, except for a few independent cases (e.g., R4 and R5 in Fig. 3B). Noticeably, unlike the very large majority of Cache domains that are located extracellularly (12), these gammaproteobacterial sCache domains are cytoplasmic sensors, further suggesting their recent emergence and neofunctionalization. In contrast, all formate sensors in alphaproteobacteria assume the sCache fold (Fig. 3C), suggesting that the major dCache-to-sCache transition in formate sensors has occurred around the separation of alphaproteobacteria and gammaproteobacteria. A few formate-responsive sCache domains found in clostridia are closely related to those in alphaproteobacteria (e.g., R6 and R7 in Fig. 3B), suggesting possible events of horizontal gene transfer. The domain composition of formate-responsive Cache domains reveals that they are almost exclusively present in chemoreceptors (Data S1).

The analysis of the isolation sites of strains harboring formate-responsive chemoreceptors shows that they are abundant in soil and freshwater (Fig. 3D). In addition, a significant number of strains were isolated from sites that are typically associated with an anaerobic metabolism (sediment, sludge, excrements), indicating the use of formate as terminal electron acceptor. Almost half of all sCache containing chemoreceptors are present in plant-associated strains or the rhizosphere, suggesting a role of formate chemotaxis in plant colonization/infection (Fig. 3D).

### Experimental verification of predicted formate-binding Cache domains

From the non-redundant set of 164 monomodular domains that were predicted to bind formate, we selected five representatives from alphaproteobacteria, gammaproteobacteria, and clostridia for experimental verification (R3 to R7 in Fig. 3B, Table 1). Four of these proteins were found to be soluble, and were subjected to microcalorimetric titrations that showed that all four proteins bound formate with *K*_D_ values ranging from 4 to 63 µM (Fig. 4, Table 1).

**Fig. 4.**
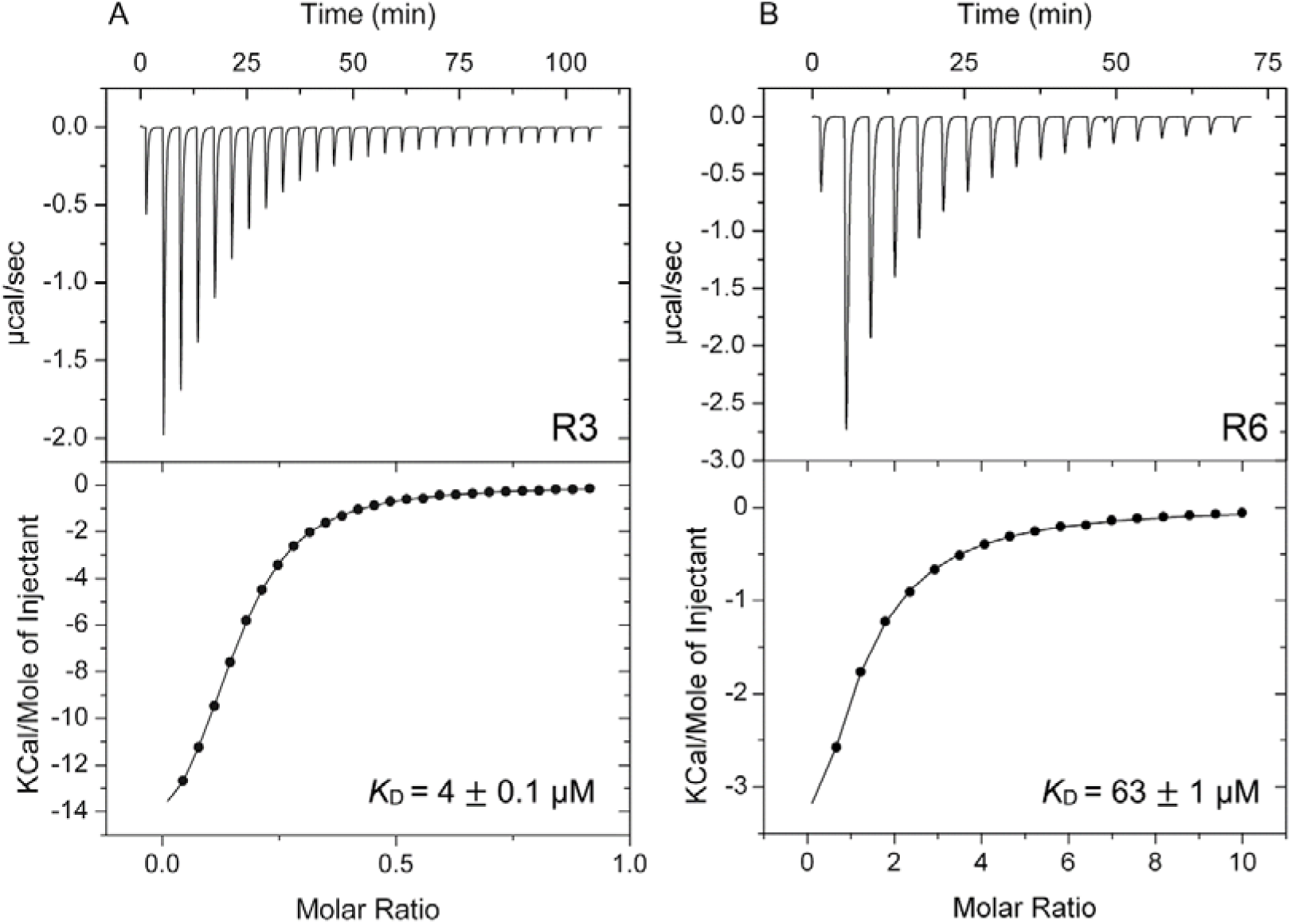
Isothermal titration calorimetry studies of selected monomodular domains with formate. **A)** Titration of 100 µM of R3 with 1 mM Na formate. **B)** Titration of 50 µM of R6 with 5 mM Na formate. Upper panels: Raw titration data. Lower panels: Integrated, concentration-normalized and dilution heat corrected raw data that were fitted with the “One-binding site” model of the MicroCal version of ORIGIN. The derived dissociation constants are provided in Table 1.

### The molecular basis for formate recognition at sCache domains

To determine the structural reasons for formate recognition we have solved the three-dimensional structures of the LBDs from chemoreceptors of *Asticcacaulis benevestitus* (R3) and *Oscillibacter ruminantium* (R6) in complex with formate to resolutions of 2.1 and 1.75 Å, respectively. Both domains assume a typical sCache fold characterized by a long N-terminal helix followed by an α/β-fold (Fig. 5 A, D). The structural alignment of both structures with all entries of the protein data bank revealed similarities primarily with the membrane proximal and distal modules of various dCache domains (Table S2).

**Fig. 5.**
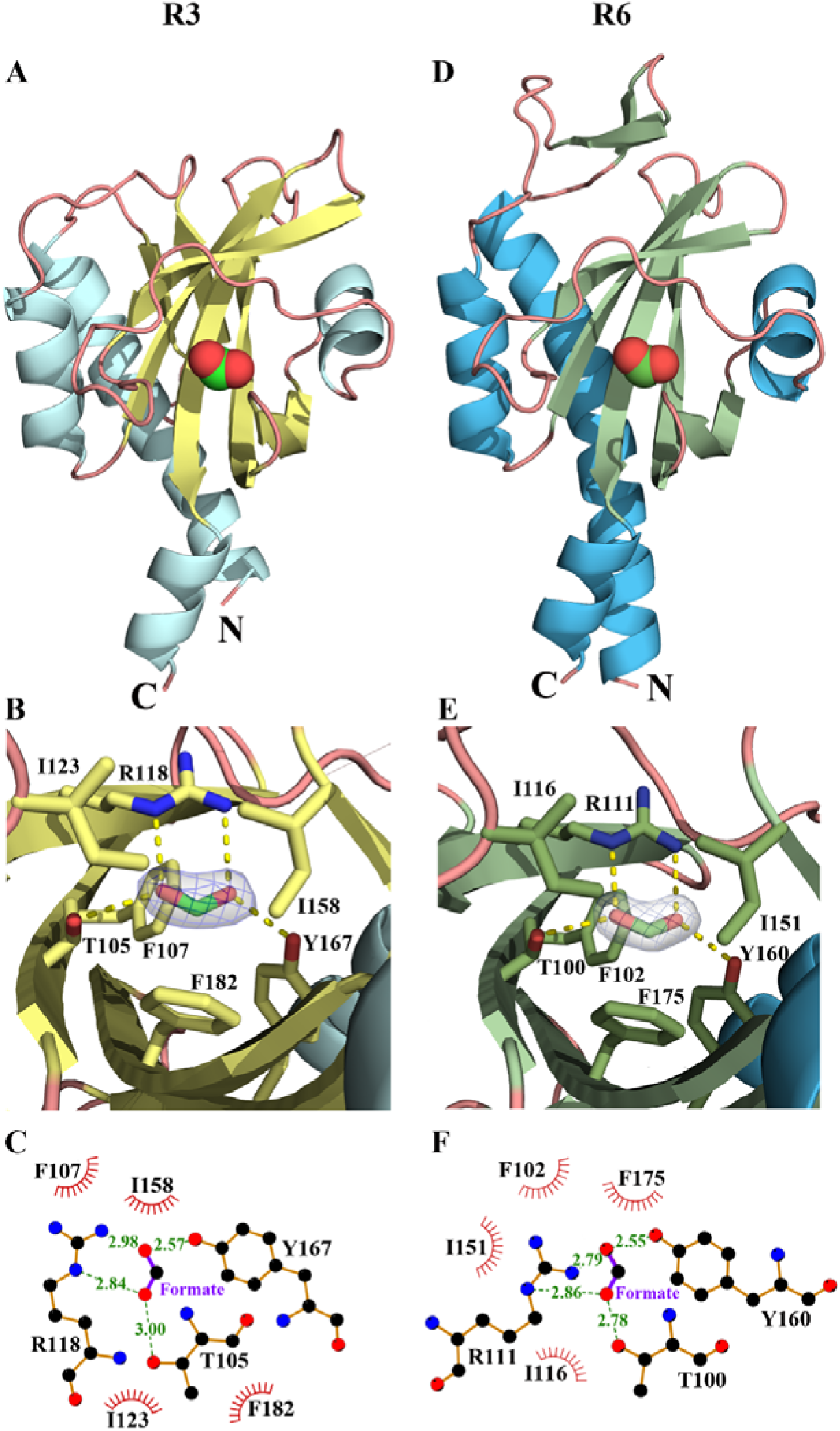
The three-dimensional structures of monomodular formate binding Cache domains. LBD structures of chemoreceptors from *Asticcacaulis benevestitus* (R3, panels A-C)) and *Oscillibacter ruminantium* (R6, panels D-F) in complex with formate. **A, D**) Overall fold with bound formate shown in spacefill mode. **B, E**) Zoom at the ligand binding site showing the |2Fo–Fc| electron density map contoured at 2.0 σ level for formate. Resides involved in formate binding are shown in stick mode. Dashed lines indicate hydrogen bonds. **C, F**) Schematic representation of residues involved in formate binding. Green numbers indicate the length of hydrogen bonds (in Å). Spoked arcs indicate hydrophobic interactions.

Well-defined electron density was observed in both structures for bound formate (Fig. 5 B, E) that permitted the precise placement of the formate model. The mode of formate recognition at R3 and R6 is almost identical. The arginine residue 118/111 (numbering for R3 and R6) plays a key role and establishes a salt bridge with formate (Fig. 5 C, F). This arginine residue corresponds to PacF R142 and Atu0526 R115, and in both cases their replacement abolished formate binding (Fig. 2D)(39). In addition to this central arginine, two hydrogen bonds are established with bound formate involving the hydroxyl groups of T105/100 and Y167/160 (Fig. 5 C, F). Furthermore, hydrophobic contacts are established involving two isoleucine and tyrosine sidechains.

R3 and R6 are from phylogenetically distant species (alphaproteobacteria and clostridia) and their LBDs share 34 % sequence identity (Table S1). However, both structures can be closely superimposed with an RMSD of Cα atoms of 0.97 Å (Fig. 6A).

**Fig. 6.**
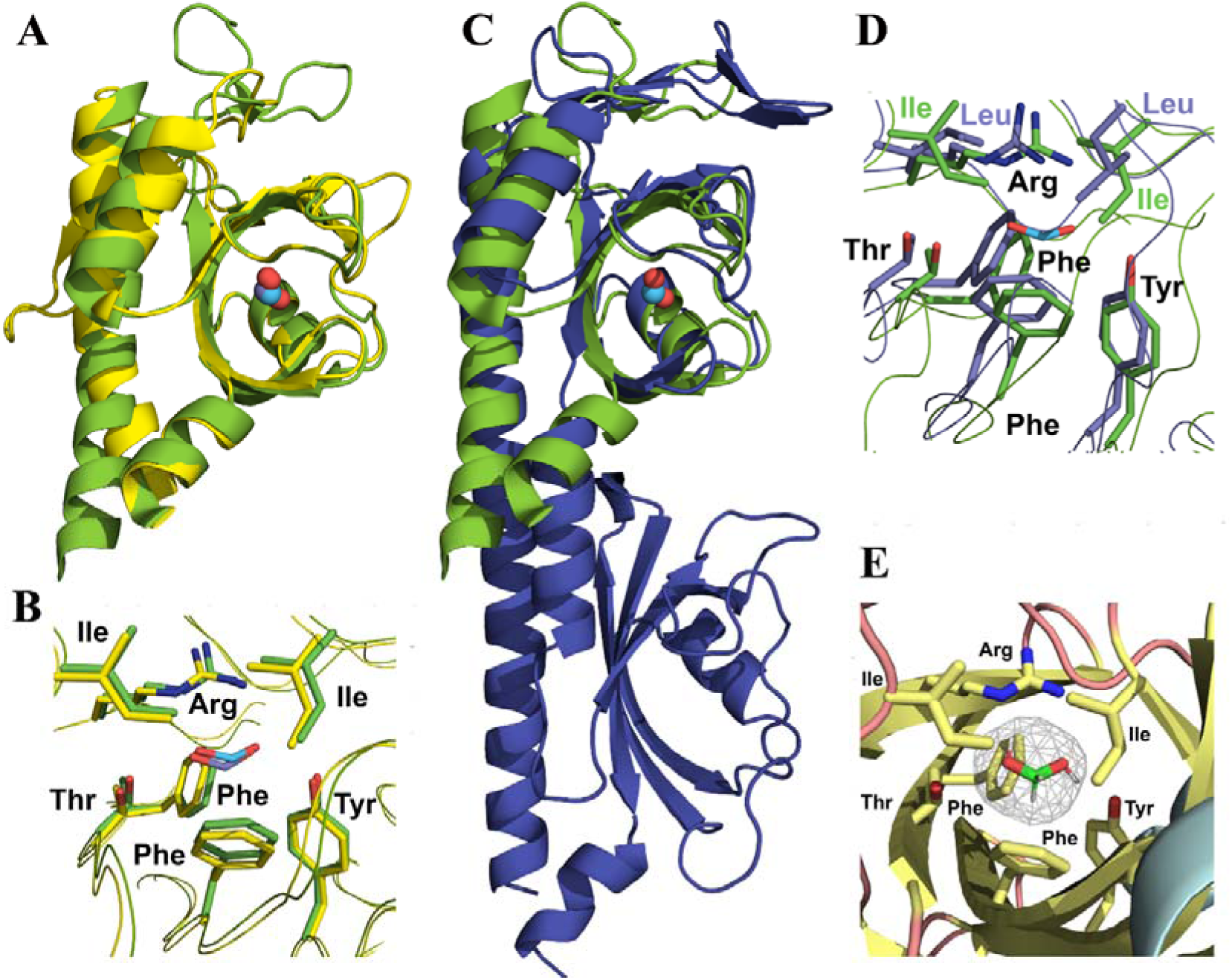
Structural features of formate recognition by mono- and bimodular domains. **A, B)** Structural alignment of the LBDs of chemoreceptors from *Asticcacaulis benevestitus* (R3, yellow)) and *Oscillibacter ruminantium* (R6, green). **A)** Overall alignment, bound formate is shown in spacefill mode. **B)** Zoom at the formate binding pockets. Amino acids involved in formate binding are labelled. **C, D)** Structural alignment of the LBD of R6 (green) with the membrane distal module of the AlphaFold2 model of the PacF LBD (violet). **C)** Overall alignment, shown is the formate molecule present in the R6 structure. **D)** Zoom at the formate binding pockets. Amino acids involved in formate binding are labelled. **E)** Zoom at the formate binding pocket of R3. The mesh shows the cavity volume as identified by Pymol using a cavity detection radius and cutoff to 3 and 5 solvent radii, respectively. Bound formate is shown in stick mode.

In this superimposition, the architecture of the binding pocket is nearly identical (Fig. 6B). As detailed above, the monomodular formate binding domains have arisen from their bimodular predecessors. The monomodular formate binding domains, as exemplified by R6 in Fig. 6C, align well with the membrane-distal module of the PacF-LBD model (RMSD of 2.58 Å). A zoom at the ligand binding pocket shows that the overall architecture and composition of binding site residues is conserved in monomodular and bimodular formate binding domains (Fig. 6D). Conserved are the arginine, the two phenylalanine, the threonine and tyrosine, whereas two PacF leucine residues are found at the position of the two isoleucine residues of R6 (Fig. 6D). This conservation pattern can be used as a marker for formate binding by uncharacterized Cache domains.

Formate is among the smallest and structurally simplest signal molecules. As detailed above, PacF ligand screening using 760 compounds resulted in the identification of a single compound, formate, indicative of high ligand specificity. What are thus the mechanisms that permit the specific recognition of such a simple molecule? The analysis of both structures indicates that ligand size exclusion by a very small binding pocket may be the primary mechanism. Using Pymol2 (42) we have determined the size of the ligand binding pocket in R3 that is shown as a mesh in Fig. 6E. The formate molecule fits closely into this space. The phenylalanine side chain at the bottom of the binding pocket (F182/175 for R3 and R6) plays a crucial role in delimiting the size of the pocket. In fact, the distances of the formate carbon atom to the atoms of the phenylalanine side chain are between 3.5 to 3.7 Å, prohibiting the binding of larger molecules, such as acetate.

## Discussion

Gene duplication is the major mode of innovation in bacterial chemoreceptors. Paralogous chemoreceptors, such as Tsr and Tar in *Escherichia coli* and *Salmonella enterica* (43), PctA, PctB, and PctC in *Pseudomonas aeruginosa* (44), and Tlp2, Tlp3, and Tlp4 in *Campylobacter jejuni* (45) provide the bacteria with a broad spectrum of sensing capabilities. Horizontal gene transfer (46), domain swap (47) and domain acquisition (48) have also been shown to drive evolution of bacterial chemoreceptors. During all these evolutionary events LBDs undergo multiple substitutions in amino acid sequence while maintaining their structural complexity. Most LBDs comprise a single structural module, however, others contain two structural modules (2) and more recently the first structure of a trimodular LBD has been reported (49). Cache domains, the predominant superfamily of extracytosolic LBDs, are present in all major families of bacterial receptors (12) and they are also identified in archaea and eukaryotes, including humans (8). Cache domains are extremely diverged and their current classification in domain databases, such as InterPro (50), includes dozens of families that belong to either a monomodular (sCache) or bimodular (dCache) structural group. Evidence was presented that strongly suggests that some bimodular dCache domains have arisen from the fusion of two monomodular sCache domains, and other bimodular dCache domains originated from duplication of monomodular sCache domains (12). In either case, novel capabilities have arisen by increasing structural complexity. Evolution by increasing complexity is generally believed to be a dominant force, however complexity can either increase or decrease during the evolution of various life forms (51). Here we present evidence that a monomodular formate-binding sCache domain has originated from a bimodular dCache domain, which binds formate only at one of its two modules, thus demonstrating that sensory domains can evolve not only by increasing, but also by decreasing complexity.

The functional advantage of dCache domains over sCache domains is unclear, because the large majority of dCache domains bind their ligands at the membrane distal module, whereas the membrane proximal module remains unoccupied (15–22). There are only a few examples of dCache domains that bind ligands at both modules (23, 24). Therefore, it can be hypothesized that the apparent lack of function may have resulted in the loss of the proximal module. This loss translates into a lower metabolic burden, because sCache domains are ∼100 amino acids shorter than dCache domains. We showed that the loss of the proximal bundle did not alter in a significant manner the mode of ligand binding (Fig. 6D) nor did it alter binding affinities (Table 1). dCache domains bind many different signal molecules (3). Therefore, this loss of complexity that we have observed for the formate-binding dCache domain may have also occurred in other bimodular domain families.

Accessing nutrients appears to be the major force that has driven the evolution of chemotaxis (52). A clear link also exists between formate chemotaxis and metabolism in *P. atrosepticum* SCRI1043. This strain harbors a functional formate hydrogenlyase-2 complex that permits the production of hydrogen from formate under anaerobic conditions and is considered a model strain for studying this class of enzymes (38). The relationship between formate chemotaxis and formate respiration is also supported by the observation of an increase in *pacF* transcript levels under anaerobic conditions (35) and the induction of formate chemotaxis when cells were grown anaerobically (Fig. 2C). The important role of formate in the anaerobic metabolism of *P. atrosepticum* SCRI1043 is supported by its failure to support aerobic growth as carbon source (Fig. S2).

There is a significant number of different formate sensing chemoreceptors. Apart from the Cache_3-Cache_2 domain containing PacF and the family of receptors with a monomodular LBD identified in this study, the Tlp1 chemoreceptor of *Campylobacter jejuni* was also found to bind specifically formate and to mediate chemoattraction (53). Although not annotated as such in public databases, its LBD appears to be a dCache-like domain according to AlphaFold modelling (40). However, dCache domains of PacF and Tlp1 employ different sensing mechanisms. While PacF binds formate at its membrane distal module, Tlp1 binds formate at its membrane proximal module (53). dCache domains of PacF and Tlp1 are distantly related as they share only 13 % sequence identity (Fig. S4). In addition, low affinity formate chemotaxis was also mediated by the sCache_2 domain containing chemoreceptor McpV of *Sinorhizobium meliloti* (54). This diversity of formate responsive chemoreceptors suggests an important physiological role of this compound.

Cache domains are universal sensing modules present in all families of bacterial receptors and are omnipresent in life. This study advances our knowledge on the evolution of this important domain family and will motivate research to establish to what degree similar evolutionary events have occurred in other domain families.

## Materials and Methods

### Strains and Plasmids

The strains, plasmids and oligonucleotides used are listed in Table S3.

### Protein overexpression and purification

All proteins were overexpressed in *E. coli* BL21(DE3) according to Rico-Jiménez et al. (55), with the exception that buffers C (30 mM Tris/HCl, 300 mM NaCl, 5 % (v/v) glycerol, 10 mM imidazole, pH 8.0) and D (20 mM Tris/HCl, 500 mM NaCl, 5 % (v/v) glycerol, 10 mM imidazole, pH 8.0) were used instead of buffers A and B, respectively. Freshly purified proteins were dialyzed into 3 mM Tris, 3 mM PIPES, 3 mM MES, 150 mM NaCl, 10 % (v/v) glycerol) at pH 8.0 (for ECA_RS17860-LBD) and at pH 6.0 (for the remaining proteins). The sequences of proteins analyzed are provided in Table S4.

### Thermal shift assay

The compound arrays PM1 and PM2A (carbon sources), PM3B (nitrogen sources), PM4A (phosphorus and sulfur sources), PM5 (nutrient supplements), PM6, PM7 and PM8 (peptide nitrogen sources) from Biolog Inc. (Hayward, CA, USA) were used. The detailed experimental protocol of the Differential Scanning Fluorimetry based ligand screening has been reported in (56). Briefly, assays were carried out using a MyIQ2 Real-Time PCR instrument (BioRad, Hercules, CA, USA). Ligand solutions were prepared by dissolving the array compounds in 50 µl of MilliQ water, which, according to the manufacturer, corresponds to a concentration of 10–20 mM. Experiments were conducted in 96-well plates and each assay mixture contained 20.5 µl of the dialyzed protein (at 10-30 µM), 2 µl of 5 X SYPRO orange (Life Technologies, Eugene, Oregon, USA) and 2.5 µl of the resuspended array compounds or the equivalent amount of buffer. Samples were heated from 23 °C to 85°C at a scan rate of 1 °C/min. The protein unfolding curves were monitored by detecting changes in SYPRO Orange fluorescence. The Tm values were determined using the first derivatives of the raw fluorescence data.

### Isothermal titration calorimetry

Experiments were conducted on a VP-microcalorimeter (Microcal, Amherst, MA, USA) at a temperature of 20 °C (PacF-LBD, PacF-LBD R142A, PacF-LBD T145A, PacF-LBD T158A) or 25 °C (remaining proteins). Freshly purified and dialyzed proteins at 9 to 100 µM were titrated with 500 µM to 5 mM ligand solutions made up in dialysis buffer. A single injection of 1.6 µl was followed by a series of 4.8 µl aliquots. The mean enthalpies from the injection of ligand solutions into the buffer were subtracted from raw titration data. Data were normalized with the ligand concentrations and fitted with the ‘One Binding Site’ model of the MicroCal version of ORIGIN (Microcal, Amherst, MA, USA).

### Quantitative capillary chemotaxis assay

Overnight cultures of *P. atrosepticum* SCRI1043 and the *pacF* mutant in minimal medium (7 g/l K_2_HPO_4,_ 2 g/l KH_2_PO_4,_ 7.5 mM (NH_4_)_2_SO_4_, 0.41 mM MgSO_4_) supplemented with 15 mM glucose were used to inoculate fresh MS medium to an OD_660_ of 0.1 (aerobic growth) or 0.15 (anaerobic growth). Under aerobic conditions cells were grown at 30 °C with shaking at 200 rpm until they reach at OD_660_ of 0.3-0.4. For anaerobic growth cells were grown at 30 °C without shaking for 5 hours in 100-ml infusion bottles under nitrogen gas. Cells were then collected by centrifugation (1,667 x *g* for 5 min at room temperature), washed gently twice with chemotaxis buffer (50 mM KH_2_PO_4_/K_2_HPO_4_, 20 mM EDTA, 0.05 % (v/v) glycerol, pH 7.0) and resuspended in the same buffer at an OD_660_ of 0.1. Aliquots (230 µl) of the resulting cell suspension were placed into the wells of 96-well microtiter plates. One μl Microcaps capillaries (Drummond Scientific, Broomall, PA, USA) were heat-sealed at one end and filled with buffer (control) or chemoeffector solution prepared in chemotaxis buffer. The capillaries were rinsed with sterile water and immersed into the bacterial suspensions at their open ends. After 30 min, capillaries were removed from the wells, rinsed with sterile water, and emptied into 1 ml of chemotaxis buffer. Serial dilutions were plated onto minimal medium plates supplemented with 20 mM glucose, incubated at 30°C prior to colony counting. Data were corrected with the number of cells that swam into buffer containing capillaries. Data are the means and standard deviations of three biological replicates conducted in triplicate.

### Generation of a ECA_RS17860 (PacF) mutant

A deletion mutant of the *pacF* gene in SCRI1043 was constructed by homologous recombination using a derivative plasmid of the suicide vector pKNG101. The plasmid was generated by amplifying the up- and downstream flanking regions of the gene to be mutated. The resulting PCR products were digested with the enzymes specified in Table S3 and ligated in a three-way set-up into pUC18Not, giving rise to plasmid pUC18Not-PacF. Subsequently, the kanamycin resistance cassette *Km3* from the plasmid p34S-km3 was inserted into the BamHI site of pUC18Not-PacF, resulting in plasmid pUC18Not-PacF-Km. The ΔECA3615-Km deletion construct was then subcloned into the marker exchange vector pKNG101 using NotI resulting in pKNG-PacF-Km. The plasmid was sequenced and carried the deletion mutant allele for the replacement of wild-type gene in the chromosome. The plasmid was then transferred into *P. atrosepticum* SCRI1043 by biparental conjugation using *E. coli* β2163. The *pacF* mutant was selected by the ability to grow on minimal medium (+ 15 mM glucose) agar plates supplemented with 50 μg/ml kanamycin and the failure to grow in presence of 50 μg/ml streptomycin. In parallel, the deletion of the gene was confirmed by PCR using DNA genomic and the primers 1F-PacF-EcoRI/2R-PacF-PstI.

### Growth experiments and determination of the minimal inhibitory concentration

Overnight cultures in minimal medium supplemented with 20 mM glucose were washed twice and diluted in fresh medium to an OD_660_ of 0.02 containing 1, 5 and 10 mM glucose or formate as sole carbon sources. Growth was monitored in an automated BioScreen C MBR instrument (Growth Curves USA, Piscataway, NJ) for 48 h using Bioscreen 100-well honeycomb microplates. Minimal inhibitory concentrations were determined by serial dilutions of formate in minimal medium cultures supplemented with 20 mM glucose. Growth was determined in a 96-well plate reader TECAN® Sunrise^TM^.

### Bioinformatics

Five thousand Atu0526 (WP_121650307.1) homologs were collected from the RefSeq database using a BLAST search (57). The sequence redundancy was reduced to 98% identity using Jalview (58) resulting in 744 non-redundant sequences that were used for analysis. Based on a multiple sequence alignment we extracted the ligand-binding modules from these proteins (e.g., the sCache domain of Atu0526-LBD and the distal module of PacF-LBD) and constructed a maximum likelihood tree using MEGA (59). The amino acid sensor PctA (NP_252999.1) is an unrelated protein from *Pseudomonas aeruginosa* (44), and the distal module of its dCache_1 was used as the outgroup to root the tree.

### Crystallization and structure resolution of the LBD of WP_040662586-LBD (R6) and WP_018081388-LBD (R3)

Formate wad added to a final concentration of 10 mM to proteins in 3 mM Tris, 3 mM PIPES, 3 mM MES, 150 mM NaCl and 10 % (v/v) glycerol, pH 6.0. Excess of formate was removed by rounds of protein concentration using 10 kDa cut-off centricon concentrators (Amicon) and subsequent dilution with the above buffer. Hanging-drop vapor diffusion and the capillary counter-diffusion trails were made using protein at 20 to 30 mg/ml. All crystallization experiments were kept at 293 K and inspected regularly. The final crystallization conditions are provided in Table S5. Data collection was done at the Xaloc beamline of the Alba Spanish synchrotron radiation source (Barcelona, Spain). Data were indexed and integrated with XDS (60) and scaled and reduced with AIMLESS (61) of the CCP4 program suite (62). Initial structural models were generated by AlphaFold2 (40) to feed Morel (63). Refinement was initiated with phenix.refine (64) of the PHENIX suite (65) and Refmac (66) of the CCP4 program suite. After manual building, ligand identification was done in Coot (67) and final water inspection and refinement was assessed including Titration-Libration-Screw parameterization (68). Towards the end of the refinement, the models were run through the PDB-REDO (69) server for verification. Both models were further verified with Molprobity (70). Table S5 summarizes X-ray data statistics and the characteristics of deposited models. Coordinates and the experimental structure factors have been deposited at the Protein Data Bank with ID 8PY0 and 8PY1.

## Acknowledgements

This study was supported by grants from the Spanish Ministry for Science and Innovation/*Agencia Estatal de Investigación* 10.13039/501100011033 (grants PID2020-112612GB-I00 to TK, PID2019-103972GA-I00 to MAM and PID2020-116261GB-I00 to JAG), the Junta de Andalucía (grant P18-FR-1621 to TK), and the NIH (grant R35GM131760 to IBZ).

## Abbreviations

HMM: hidden Markov model
LBD: ligand binding domain

