## Supplementary Figures and Tables for "Bacterial sensor evolved by decreasing complexity"

to

by

Elizabet Monteagudo-Cascales, José A. Gavira, Jiawei Xing, Félix Velando, Miguel A. Matilla, Igor B. Zhulin and Tino Krell

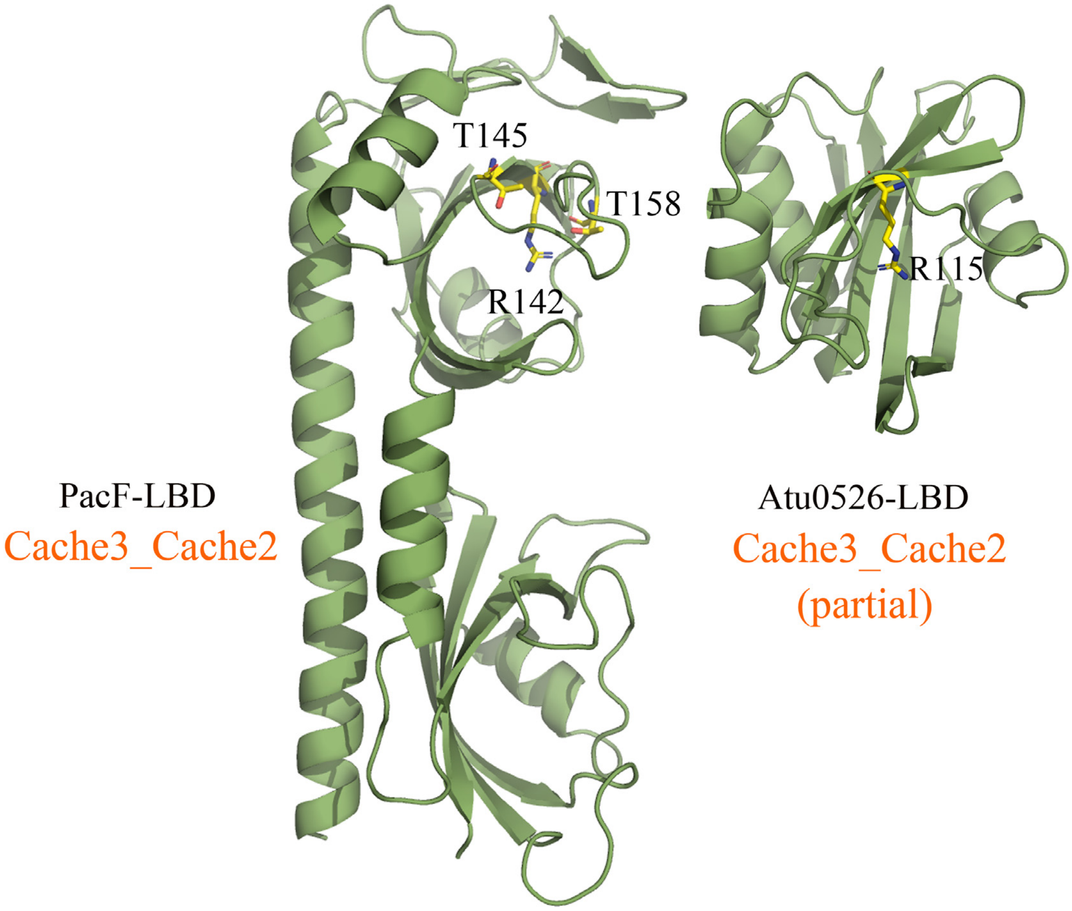


**Fig. S1) Alphafold models of PacF-LB and Atu0526-LBD.** Amino acids involved in formate binding are shown in stick mode. The annotation of domains in the Pfam database is shown in orange.

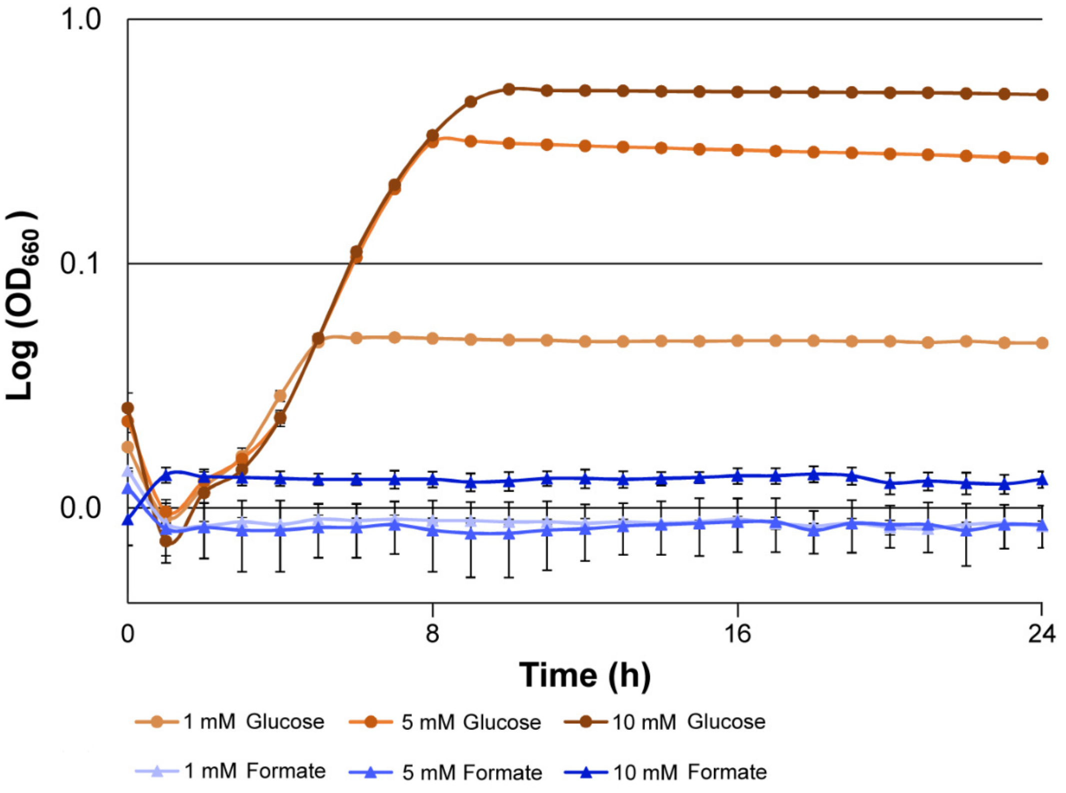


**Fig. S2) Aerobic growth experiments of *P. atrosepticum* SCRI1043 in minimal medium supplemented with different concentrations of glucose (control) and formate.** Shown are means and standard deviations from three replicates. Standard deviation for experiments with glucose are smaller than the size of the data points.

```
                   40        50        60        70        80        90
                    |         |         |         |         |         |
PacF          LTRSAGDQLKALTLNDMQSQVNGVTDMINMYDTSLQAEVSNYTRLLASFLPTQFSLDTVN
Atu0526       ---RLKDNIITTALRDTHGAMRG---MAILYEMKVGGVALEMV---------DGDLKSVG
R6            QSFRVTNNIVDMQVQDTLKGAAN---MLGLYLEEQFGPLSLNV---------AGNLVDVD
R3            -VVQMRADTASTALKYQHSALRVA--SATLHRQFPDTSVEWAP---------DGNVQKVV

                   100       110       120       130         140       150
                    |         |         |         |           |         |
PacF          RTPFSNTSHPTLKAGDTVLNLNTDVTDDFLSRTHAISTIFVRD--GEDFTRITTSLKKED
Atu0526       RASIGTM-------------RDNDLVDRTAAGNGGIATVFEAK--AGEFIRLTTNLKNEK
R6            GRPIE---------------GENDYIDRLSQSMNVVATVFAKN--GNDYIRTLTTIKDDN
R3            MDTVPTF-------------TDHAMIDEIARVSGQQATLFAFDPAQDDFIRTTTSITKPD

                     160       170       180       190       200       210
                      |         |         |         |         |         |
PacF          GSRAMGTKLDRESPAYALVIKGETYSGLATLFGKQYITQYQPIHDGENKVIGILFVGVDI
Atu0526       GERAAGTKLATDHPAFEKVSKGEAYFGTATLFGTSYMTGYMPVTNKTGATVGILFVGVPM
R6            GERVVGTALDSSGDAYRTLNAGGTYFGEATILGSAYMTGYVPLLDRTGQAIGACFVGVSI
R3            GSRAVGTNLGQDSKAFAPIKAGKTYLGKADILGTSYYTIYAPVFNTRGDVTGILFSGVKT

                     220       230       240       250       260       270
                      |         |         |         |         |         |
PacF          TKQFTEMQQTILNKKMGETGHFFVLSAKKGKDAGNYLYHQSNANQRPDWSEGALEKVLAT
Atu0526       DFYNAQIYSLRDM-----------------------------------------------
R6            ESVNAILNEGMASAIRS-------------------------------------------
R3            ATVQEAANALMMK-----------------------------------------------

                     280       290       300       310       320
                      |         |         |         |         |
PacF          GNGTIEYDNNTMKGEEKTRIMVYRSIPQWNWIVAGTISKESLMAEVNRTRNLFL
Atu0526       ------------------------------------------------------
R6            ------------------------------------------------------
R3            ------------------------------------------------------
```

**Fig. S3) Sequence alignment of the LBDs of formate-responsive sensor domains.** PacF residues that have been replaced by alanine (Table 2) are shaded in cyan. Residues that were shown to be involved in signal binding are shaded in yellow. The numbering corresponds to that of the full-length PacF. The alignment was done using the CLUSTALW algorithm of the npsa suite (1) using the GONNET weight matrix, a gap opening penalty of 10 and a gap extension penalty of 0.2. red: identical; green: highly similar; blue: weakly similar.

```
                           40        50        60        70        80
                           |         |         |         |         |
PacF_LBD      -------LTRSAGDQLKALTLNDMQSQVNGVTDMINMYDTSLQAEVSNYTRLLASFLPTQ
Tlp1_LBD      TKQVSQNITKNTEDILASITK-EYATQTQGIFGEMIALNKSISGTLTEMFRSTSKEDLDI

                 90       100       110         120       130       140
                 |         |         |           |         |         |
PacF_LBD      FSLDTVNRTPFSNTSHPTLKAGDTVLN--LNTDVTDDFLSRTHAISTIFVRDGEDFTRIT
Tlp1_LBD      DNITNIITNTFDNSAYSNFTYLYLIDPPEYFKEESKFFNTQSGKFVMLYADEEKDNKGGI

                  150          160       170       180       190       200
                   |            |         |         |         |         |
PacF_LBD      TSLKKEDGSR---AMGTKLDRESPAYALVIKGETYSGLATLFGKQYITQYQPIHDGENKV
Tlp1_LBD      KAIQASDEIANLQVVQDILKKAKYGENKVYIGRPIKMNLEGQDFDAVNVAIPIFDRKNQV

                     210       220       230       240       250       260
                      |         |         |         |         |         |
PacF_LBD      IGILFVGVDITKQFTEMQQTILNKKMGETGHFFVLSAKKGKDAGNYLYHQSNANQRPDWS
Tlp1_LBD      VGVIGMTLDFSDIATYLLDPKGQKYDGELRVLLNSDGFMAIHPNKNLVLKNLKDINPNKG

                     270        280       290       300       310       320
                      |          |         |         |         |         |
PacF_LBD      EGALEKVLATG-NGTIEYDNNTMKGEEKTRIMVYRSIPQWNWIVAGTISKESLMAEVNRT
Tlp1_LBD      AQETYKAISEGKNGVFNYIAS--DGDDSYAAINSFKVQDSSWAVLVTAPKYSVFKPLKKL


PacF_LBD      RNLFL
Tlp1_LBD      QL---
```

**Fig. S4) Sequence alignment of the LBDs of the PacF and Tlp1 formate chemoreceptors from *Pectobacterium atrosepcticum* SCRI1043 and *Campylobacter jejuni* NCTC 11168, respectively**. The numbering corresponds to that of full-length PacF. The alignment was done using the CLUSTALW algorithm of the npsa suite (1) using the GONNET weight matrix, a gap opening penalty of 10 and a gap extension penalty of 0.2. red: identical; green: highly similar; blue: weakly similar.

**Table S1) Sequence identities of formate-responsive LBDs.** Values were obtained using VectorBuilder (https://en.vectorbuilder.com/).

| | PacF[a] | Atu0526 | R3 | R6 |
|---|---|---|---|---|
| PacF | X | 36 | 34 | 32 |
| Atu0526 | | X | 36 | 40 |
| R3 | | | X | 34 |
| R6 | | | | X |

[a]Identity of aligned sections is shown, excluding the membrane-proximal module of PacF-LBD.

**Table S2) Structural alignment of the LBDs of receptors R3 and R6 with all structures deposited in the protein data bank. Listed are the structures with the best alignment.**

| Pdb ID | Name | Sensor protein family[a] | Sensor domain family/module | ligands | Bacterial species | Z-score | rmsd | lali | Ident (%) | Ref. |
|---|---|---|---|---|---|---|---|---|---|---|
| **R3** | | | | | | | | | | |
| 6lnp | Citrate biosensor based on CitA | Synthetic construct | sCache | citrate | *Klebsiella pneumoniae* | 11.9 | 2.7 | 123 | 14 | (2) |
| 3lif | rpHK1S-Z16 | DGC | dCache-P[b] | - | *Rhodopseudomonas palustris* | 10.6 | 3.3 | 116 | 9 | (3) |
| 6fu4 | TlpQ | CR | dCache-P | histamine | *Pseudomonas aeruginosa* | 10.5 | 3.5 | 124 | 15 | (4) |
| 7prq | PctD | CR | dCache-P | choline | *Pseudomonas aeruginosa* | 10.5 | 3.2 | 126 | 11 | (5) |
| 5wbf | TlpC | CR | dCache-P | lactate | *Helicobacter pylori* | 10.5 | 2.7 | 118 | 11 | (6) |
| 3by9 | DctB | HK | dCache-D | succinate | *Vibrio cholerae* | 10.3 | 2.8 | 125 | 14 | (7) |
| 2zbb | DctB | HK | dCache | malonic acid | *Escherichia coli* | 10.3 | 2.8 | 123 | 11 | unpublished |
| 4jgo | KinD | HK | dCache-D | pyruvate | *Bacillus subtilis* | 10.3 | 3.0 | 122 | 13 | (8) |
| 5ere | - | unknown | Periplasmic binding protein | - | *Desulfohalobium retbaense* | 10.1 | 3.1 | 124 | 10 | unpublished |
| 5hl6 | - | unknown | GAF | - | *Burkholderia vietnamiensis* | 9.8 | 3.0 | 103 | 10 | unpublished |
| 6e0a | TlpA | CR | dCache-P | - | *Helicobacter pylori* | 9.7 | 2.9 | 115 | 6 | (9) |
| 1ojg | DucS | HK | sCache | - | *Escherichia coli* | 9.6 | 3.3 | 124 | 11 | (10) |
| 3lic | soHK1S-Z6 | HK | dCache-D | - | *Shewanella oneidensis* | 9.6 | 3.4 | 129 | 7 | (3) |
| 4ywz | WalK | HK | PAS | | *Staphylococcus aureus* | 9.5 | 3.3 | 121 | 12 | (11) |
| 6mab | RsbU | SpoIIE | dCache-D | - | *Chlamydia trachomatis* | 9.5 | 3.2 | 123 | 14 | (12) |
| 3ci6 | - | PP | GAF | - | *Acinetobacter baylyi* | 9.5 | 2.6 | 102 | 13 | unpublished |
| 6pzj | - | CR | dCache-P | - | *Leptospira interrogans* | 9.5 | 3.2 | 115 | 10 | unpublished |
| 3dte | - | TR | - | - | *Deinococcus deserti* | 9.2 | 2.5 | 101 | 12 | (13) |
| 3cwf | - | HK | sCache | - | *Bacillus subtilis* | 9.2 | 2.6 | 97 | 11 | (14) |
| 7psg | PacA | CR | dCache-P | betaine | *Pectobacterium atrosepticum* | 9.1 | 3.7 | 115 | 14 | (5) |
| **R6** | | | | | | | | | | |
| 3lif | rpHK1S-Z16 | DGC | dCache-P | - | *Rhodopseudomonas palustris* | 12.5 | 2.6 | 125 | 8 | (3) |
| 7prq | PctD | CR | dCache-P | choline | *Pseudomonas aeruginosa* | 12.4 | 3.3 | 141 | 10 | (5) |
| 4wy9 | Tlp1 | CR | dCache-P | - | *Campylobacter jejuni* | 12.2 | 2.9 | 133 | 14 | (15) |
| 4ywz | WalK | HK | PAS | | *Staphylococcus aureus* | 12.2 | 3.2 | 135 | 13 | (11) |
| 3by9 | DctB | HK | dCache-D | succinate | *Vibrio cholerae* | 12.1 | 2.8 | 134 | 10 | (7) |
| 2zbb | DctB | HK | dCache-D | malonate | *Escherichia coli* | 11.9 | 2.7 | 135 | 13 | unpublished |
| 5wbf | TlpC | CR | dCache-P | lactate | *Helicobacter pylori* | 11.7 | 3.0 | 134 | 11 | (6) |
| 6d8v | McpX | CR | dCache-D | 1,1-dimethyl-prolinium | *Sinorhizobium meliloti* | 11.6 | 3.1 | 137 | 13 | (16) |

| | | | | | | | | | | |
|---|---|---|---|---|---|---|---|---|---|---|
| 4jgo | KinD | HK | dCache-D | pyruvate | *Bacillus subtilis* | 11.5 | 2.5 | 129 | 12 | (8) |
| 5ere | - | unknown | Periplasmic binding protein | - | *Desulfohalobium retbaense* | 11.5 | 2.7 | 132 | 11 | unpublished |
| 6e0a | TlpA | CR | dCache-P | - | *Helicobacter pylori* | 11.2 | 2.8 | 132 | 8 | (9) |
| 3lic | soHK1S-Z6 | HK | dCache-D | - | *Shewanella oneidensis* | 11.1 | 3.3 | 138 | 9 | (3) |
| 5ltx | PctA | CR | dCache-P | L-Met | *Pseudomonas aeruginosa* | 10.9 | 3.2 | 131 | 14 | (17) |
| 6fu4 | TlpQ | CR | dCache-P | histamine | *Pseudomonas aeruginosa* | 10.7 | 3.6 | 133 | 11 | (4) |
| 6ior | Mlp24 | CR | dCache-D | L-Asn | *Vibrio cholerae* | 10.7 | 3.2 | 128 | 11 | (18) |
| 3lid | vpHK1S-Z8 | DGC | dCache-D | - | *Vibrio parahaemolyticus* | 10.5 | 2.8 | 136 | 10 | (3) |
| 6mni | PscC | CR | dCache-D | proline | *Pseudomonas syringae* | 10.4 | 3.1 | 130 | 11 | (19) |
| 3lib | mmHK1S-Z3 | HK | dCache-P | - | *Methanosarcina mazei* | 10.3 | 3.3 | 129 | 12 | (3) |
| 3ub9 | TlpB | CR | sCache | hydroxyurea | *Helicobacter pylori* | 10.2 | 3.5 | 131 | 9 | (20) |
| 6pzj | - | CR | dCache-P | - | *Leptospira interrogans* | 10.1 | 2.8 | 118 | 9 | unpublished |

[a] DGC, diguanylate cyclase; CR, chemoreceptor; HK, histidine kinase; SpoIIE: Stage II sporulation protein E; PP, protein phosphatase; TR, transcriptional regulator

[b] structural alignment with the membrane distal (D) or the membrane-proximal (P) module of the dCache domain.

**Table S3) Strains, plasmids and oligonucleotides used in this study.**

| **Strains** | Genotype or relevant characteristics | Reference |
|---|---|---|
| *Escherichia coli* BL21(DE3) | F⁻ *ompT gal dcm lon hsdS*$_B$($r_B^-m_B^-$) λ(DE3 [*lacI lacUV5-T7p07 ind1 sam7 nin5*]) [*malB*$^+$]$_{K-12}$(λ$^S$) | (21) |
| *E. coli* DH5α | F⁻ *endA1 glnV44 thi-1 recA1 relA1 gyrA96 deoR nupG purB20* φ80d*lacZ*ΔM15 Δ(*lacZYA-argF*)U169, hsdR17($r_K^-m_K^+$), λ⁻ | (22) |
| *E. coli CC118λpir* | *araD Δ(ara, leu) ΔlacZ74 phoA20 galK thi-1 rspE rpoB argE recA1 λpir* | (23) |
| E. coli β2163 | F⁻RP4-2-Tc::Mu Δ*dapA*::(*erm-pir*); Km$^R$, Em$^R$ | (24) |
| *Pectobacterium atrosepticum* SCRI1043 | Wild type, plant pathogen | (25) |
| *P. atrosepticum* SCRI1043-PacF | Deletion mutant of *ECA_RS17860*, Km$^R$ | This study |
| **Plasmids** | | |
| pET28b(+) | Protein expression plasmid; Km$^R$ | Novagen |
| pET28b-PacF-LBD | Km$^R$; pET28b(+) derivative containing a DNA fragment encoding the ECA_RS17860-LBD | This study[1] |
| pET28b-PacF-LBD _R142A | KmR; pET28b(+) derivative containing a DNA fragment encoding the ECA_RS17860-LBD_R142A | This study[1] |
| pET28b-PacF-LBD _T145A | KmR; pET28b(+) derivative containing a DNA fragment encoding the ECA_RS17860-LBD_T145A | This study[1] |
| pET28b-PacF-LBD _T158A | KmR; pET28b(+) derivative containing a DNA fragment encoding the ECA_RS17860-LBD_T158A | This study[1] |
| pET28b-WP_018081388-LBD | Km$^R$; pET28b(+) derivative containing a DNA fragment encoding the WP_018081388-LBD | This study[1] |
| pET28b-WP_040662586-LBD | Km$^R$; pET28b(+) derivative containing a DNA fragment encoding the WP_040662586-LBD | This study[1] |
| pET28b-WP_134194227-LBD | Km$^R$; pET28b(+) derivative containing a DNA fragment encoding the WP_134194227-LBD | This study[1] |
| pET28b-WP_105260142-LBD | Ap$^R$, pET28b(+) derivative containing a DNA fragment encoding the WP_105260142-LBD | This study[1] |
| pET28b-WP_106058630-LBD | Km$^R$; pET28b(+) derivative containing a DNA fragment encoding the WP_106058630-LBD | This study[1] |
| pUC18Not | Ap$^R$; identical to pUC18 but with two NotI sites flanking pUC18 polylinker | (26) |
| pUC18Not-PacF | Ap$^R$; 2.0-kb PCR fragment containing the *pacF* gene cloned into the NdeI/PstI sites of pUC18Not | This study |
| pUC18Not-PacF-Km | Km$^R$, Ap$^R$; a *Km3* antibiotic cassette from p34S-Km3 cloned into BamHI site of pUC18Not-PacF | This study |
| p34S-Km3 | Km$^R$, Ap$^R$; *Km3* antibiotic cassette | (27) |
| pKNG101 | Sm$^R$; *oriR6K mob sacBR* | (25) |
| pKNG-PacF-Km | Sm$^R$, Km$^R$; 3.3 kb NotI fragment of pUC18Not-PacF-Km was cloned at the same site in pKNG101 | This study |

**Oligonucleotides**

| **Name** | **Sequence (5'-3')** | **Purpose** |
|---|---|---|
| 1F-PacF-EcoRI | TAATGAATTCACGTCAGATTGGCGAGTGTA | pUC18Not-PacF |
| 1R-PacF-BamHI | TAATGGATCCAACAATATCGCCACGCTCAG | |
| 2F-PacF-BamHI | TAATGGATCCGCGCAGAGTCTACGTCAGAA | |
| 2R-PacF-PstI | TAATCTGCAGCAATCGTTGCCGATGTCACC | |

[1]Plasmids were synthesized by GenScript (Piscataway, New Jersey, USA)

**Table S4) Sequences of proteins used in this study. The N-terminal extension containing the histidine tag is shown in bold face.**

| Name | Strain/species | Sequence |
|---|---|---|
| PacF-LBD (ECA_RS17860-LBD) | *Pectobacterium atrosepticum* SCRI10 | **MGSSHHHHHHSSGLVPRGSHM**LTRSAGDQLKALTLNDMQSQ VNGVTDMINMYDTSLQAEVSNYTRLLASFLPTQFSLDTVNR TPFSNTSHPTLKAGDTVLNLNTDVTDDFLSRTHAISTIFVR DGEDFTRITTSLKKEDGSRAMGTKLDRESPAYALVIKGETY SGLATLFGKQYITQYQPIHDGENKVIGILFVGVDITKQFTE MQQTILNKKMGETGHFFVLSAKKGKDAGNYLYHQSNANQRP DWSEGALEKVLATGNGTIEYDNNTMKGEEKTRIMVYRSIPQ WNWIVAGTISKESLMAEVNRTRNLF |
| PacF-LBD R142A | *P. atrosepticum* SCRI1043 | **MGSSHHHHHHSSGLVPRGSHM**LTRSAGDQLKALTLNDMQSQ VNGVTDMINMYDTSLQAEVSNYTRLLASFLPTQFSLDTVNR TPFSNTSHPTLKAGDTVLNLNTDVTDDFLSRTHAISTIFVR DGEDFTAITTSLKKEDGSRAMGTKLDRESPAYALVIKGETY SGLATLFGKQYITQYQPIHDGENKVIGILFVGVDITKQFTE MQQTILNKKMGETGHFFVLSAKKGKDAGNYLYHQSNANQRP DWSEGALEKVLATGNGTIEYDNNTMKGEEKTRIMVYRSIPQ WNWIVAGTISKESLMAEVNRTRNLF |
| PacF-LBD T145A | *P. atrosepticum* SCRI1043 | **MGSSHHHHHHSSGLVPRGSHM**LTRSAGDQLKALTLNDMQSQ VNGVTDMINMYDTSLQAEVSNYTRLLASFLPTQFSLDTVNR TPFSNTSHPTLKAGDTVLNLNTDVTDDFLSRTHAISTIFVR DGEDFTRITASLKKEDGSRAMGTKLDRESPAYALVIKGETY SGLATLFGKQYITQYQPIHDGENKVIGILFVGVDITKQFTE MQQTILNKKMGETGHFFVLSAKKGKDAGNYLYHQSNANQRP DWSEGALEKVLATGNGTIEYDNNTMKGEEKTRIMVYRSIPQ WNWIVAGTISKESLMAEVNRTRNLF |
| PacF-LBD T158A | *P. atrosepticum* SCRI1043 | **MGSSHHHHHHSSGLVPRGSHM**LTRSAGDQLKALTLNDMQSQ VNGVTDMINMYDTSLQAEVSNYTRLLASFLPTQFSLDTVNR TPFSNTSHPTLKAGDTVLNLNTDVTDDFLSRTHAISTIFVR DGEDFTRITTSLKKEDGSRAMGAKLDRESPAYALVIKGETY SGLATLFGKQYITQYQPIHDGENKVIGILFVGVDITKQFTE MQQTILNKKMGETGHFFVLSAKKGKDAGNYLYHQSNANQRP DWSEGALEKVLATGNGTIEYDNNTMKGEEKTRIMVYRSIPQ WNWIVAGTISKESLMAEVNRTRNLF |
| WP_040662586-LBD | *Oscillibacter ruminantium* | **MGSSHHHHHHSSGLVPRGSHM**QSFRVTNNIVDMQVQDTLKG AANMLGLYLEEQFGPLSLNVAGNLVDVDGRPIEGENDYIDR LSQSMNVVATVFAKNGNDYIRTLTTIKDDNGERVVGTALDS SGDAYRTLNAGGTYFGEATILGSAYMTGYVPLLDRTGQAIG ACFVGVSIESVNAILNEGMASAIR |
| WP_018081388-LBD | *Asticcacaulis benevestitus* | **MGSSHHHHHHSSGLVPRGSHM**QMRADTASTALKYQHSALRV ASATLHRQFPDTSVEWAPDGNVQKVVMDTVPTFTDHAMIDE IARVSGQQATLFAFDPAQDDFIRTTTSITKPDGSRAVGTNL GQDSKAFAPIKAGKTYLGKADILGTSYYTIYAPVFNTRGDV TGILFSGVKTATVQEAANA |
| WP_105260142-LBD | *Rhodoferax* sp. TS-BS-61-7 | **MGSSHHHHHHSSGLVPRGSHM**ARRVNAAARKFAASLPGVFS LDGKQRVLVQDRQVAVLRNGQHTLNLDFALPDGFSQATGLV ATVFVRDGDDFIRITTSVRKQDGQRAIGTPLDRSQPAYNDL LQGRAHLGYATIFGKQYLTRYEPVHDASGRVIGILFVGLDI TASPG |
| WP_134194227-LBD | *Paraburkholderia rhizosphaerae* | **MGSSHHHHHHSSGLVPRGSHM**PLRVQVGRHAKTLAGYFPAP FIRDESNGIAIHGAVTPRLLCGNALLNLNYAEVDRFTHATR STATLFVIKGEDFVRVTTSVKKQNGERAVGTQLDRSHPAWR LLRDGQTYTGYATLFGKQYMTQYEPIRDSAGRVIGALYVGL DVSEAWTLS |
| WP_106058630-LBD | *Clostridium vincentii* | **MGSSHHHHHHSSGLVPRGSHM**KGSVYMKEISNHHINNKLSS DINALSTYSELIYGNLTISNGNLVDENAGTIKGNYTAVDRI SKDLGDLATIFVTDGDDFIRVSTNIMDENGLRAEGTKLDTN |

| | | |
|---|---|---|
| | | SEAYKSLSKGERYIGSSTIFNVDYETVYENILDSKGDVIGA<br>YFIGIPTTTANEIITISLNSLRN |

**Table S5) Data collection and refinement statistics. Statistics for the highest-resolution shell are shown in parentheses.**

| | **R3** | **R6** |
|---|---|---|
| Source | *Asticcacaulis benevestitus* | *Oscillibacter ruminantium* |
| Crystallization conditions | 0.1 M Tris/HCl, 2.0 M $(NH_4)_2SO_4$, pH 8.5 | 10 % (w/v) PEG 8000, 100 mM $KH_2PO_4$/$Na_2HPO_4$ 200 mM NaCl, pH 6.2 |
| **Data collection** | | |
| PSB ID | 8PY1 | 8PY0 |
| Beam Line | XALOC (ALBA) | XALOC (ALBA) |
| Resolution range (Å) | 73.38 - 2.1 (2.175 - 2.10) | 50.56 - 1.75 (1.813 - 1.75) |
| Space group | I 41 2 2 | P 31 2 1 |
| Unit cell | | |
| a, b, c (Å) | 103.768 103.768 220.404 | 58.379 58.379 156.462 |
| α, β, γ (°) | 90 90 90 | 90 90 120 |
| Unique reflections | 35,374 (3397) | 31,951 (3178) |
| Multiplicity | 10.3 (5.4) | 4.8 (4.6) |
| Completeness (%) | 99.04 (96.91) | 96.81 (99.81) |
| Mean I/sigma(I) | 15.33 (2.48) | 15.57 (3.21) |
| Wilson B-factor | 33.39 | 19.67 |
| R-merge | 0.1162 (1.179) | 0.05408 (0.3324) |
| CC1/2 | 0.997 (0.701) | 0.999 (0.935) |
| CC* | 0.999 (0.908) | 1 (0.983) |
| **Refinement** | | |
| R-work/ R-free (%) | 26.65 / 30.72 | 18.82 / 21.65 |
| Number of atoms | | |
| Protein | 3583 | 2575 |
| Ligands | 19 | 8 |
| Solvent | 105 | 250 |
| Bond lengths (Å) | 0.003 | 0.004 |
| Bond angles (°) | 0.59 | 0.71 |
| Ramachandran (%) | | |
| favored | 98.03 | 98.38 |
| allowed | 1.75 | 0.97 |
| outliers | 0.22 | 0.65 |
| B-factor ($Å^2$) | 49.98 | 28.66 |
